# Generation of highly diverse peptide library by linear-double-stranded DNA based AND gate genetic circuit in mammalian cells

**DOI:** 10.1101/392498

**Authors:** Shuai Li, Weijun Su, Chunze Zhang

**Author notes:** Shuai Li and Weijun Su contributed equally to this work. To whom correspondence should be addressed.; &.

## Abstract

DNA-encoded peptide libraries are ideal functional peptide discovery platforms for their extremely large capacity. However, it’s still difficult to build high content peptide library in intact mammalian cells, which offer advantages associated with appropriate protein modification, proper protein folding, and natural status of membrane protein. Our previous work established linear-double-stranded DNAs (ldsDNAs) as innovative biological parts to implement AND gate genetic circuits in mammalian cell line. In the current study, we employ ldsDNA with terminal NNK degenerate codons as AND gate input to build highly diverse peptide library in mammalian cells. This <u>**l**</u>dsDNA-<u>**b**</u>ased <u>**A**</u>ND <u>**g**</u>ate (LBAG) peptide strategy is easy to conduct, only PCR reaction and cell transfection experiments are needed. High-throughput sequencing (HTS) results reveal that our new LBAG strategy could generate peptide library with both amino acid sequence and peptide length diversities. Moreover, by a mammalian cell two-hybrid system, we pan an MDM2 protein interacting peptide through the LBAG peptide library. Our work establishes ldsDNA as biological parts for building highly diverse peptide library in mammalian cells.

## Introduction

Different state-of-the-art display technologies, e.g., phage, ribosome, mRNA, bacterial, and yeast-display, have been developed to identify functional peptides (1–7). Through construction and screen vast peptide libraries, these display technologies physically couple between phenotype (high affinity high selectivity binding) and genotype (DNA sequence). Currently, plasmids and PCR products have been explored as genetic mediums to store highly diverse coding information of the peptide library (8,9). These genetic materials are introduced into phage-host bacteria, yeast, or test tubes to transcript-translate into peptides. However, it’s still a big challenge to build high content peptide library in intact mammalian cells. Some attempts have been made to introduce DNA-encoded peptide library into mammalian cells using episomal-, viral- or transposon-mediated gene transfer (10–17). While, the peptide library capacity is constantly lost during plasmid extraction, lentivirus generation, transfection/infection process. This leads to small library size relative to other display technologies.

We previously applied linear-double-stranded DNA (ldsDNA, or PCR amplicon) as novel biological parts to implement Boolean logic AND gate genetic circuits in mammalian cells (18). Via splitting essential gene expression cassette into signal-mute ldsDNA, our strategy achieves two or three-input AND gate calculation with a low noise-signal ratio both *in vitro* and *in vivo*. This split-relink process is similar to V(D)J recombination, which serves as the fundamental molecular mechanism to generate highly diverse repertoires of antigen-receptors (antibodies and T-cell receptors) in mammalian (19,20). In the present study, we employ this <u>**l**</u>dsDNA-<u>**b**</u>ased <u>**A**</u>ND-<u>**g**</u>ate (LBAG) genetic circuit to generate high content peptide library.

## Results

### Generation of peptide library by ldsDNA-based AND-gate genetic circuit

Our previous study demonstrated that ldsDNAs (PCR amplicons), split from intact gene expression cassettes (containing promoter, gene coding sequence and polyA signal), could undergo reconnection to form one output-signal-generating molecule in cultured cells (18). Here we hypothesize that ldsDNAs with terminal NNK degenerate codons could generate highly diverse peptide library through <u>**l**</u>dsDNA-<u>**b**</u>ased <u>**A**</u>ND-<u>**g**</u>ate (LBAG) calculation in mammalian cells (Figure 1).

**Figure 1.**
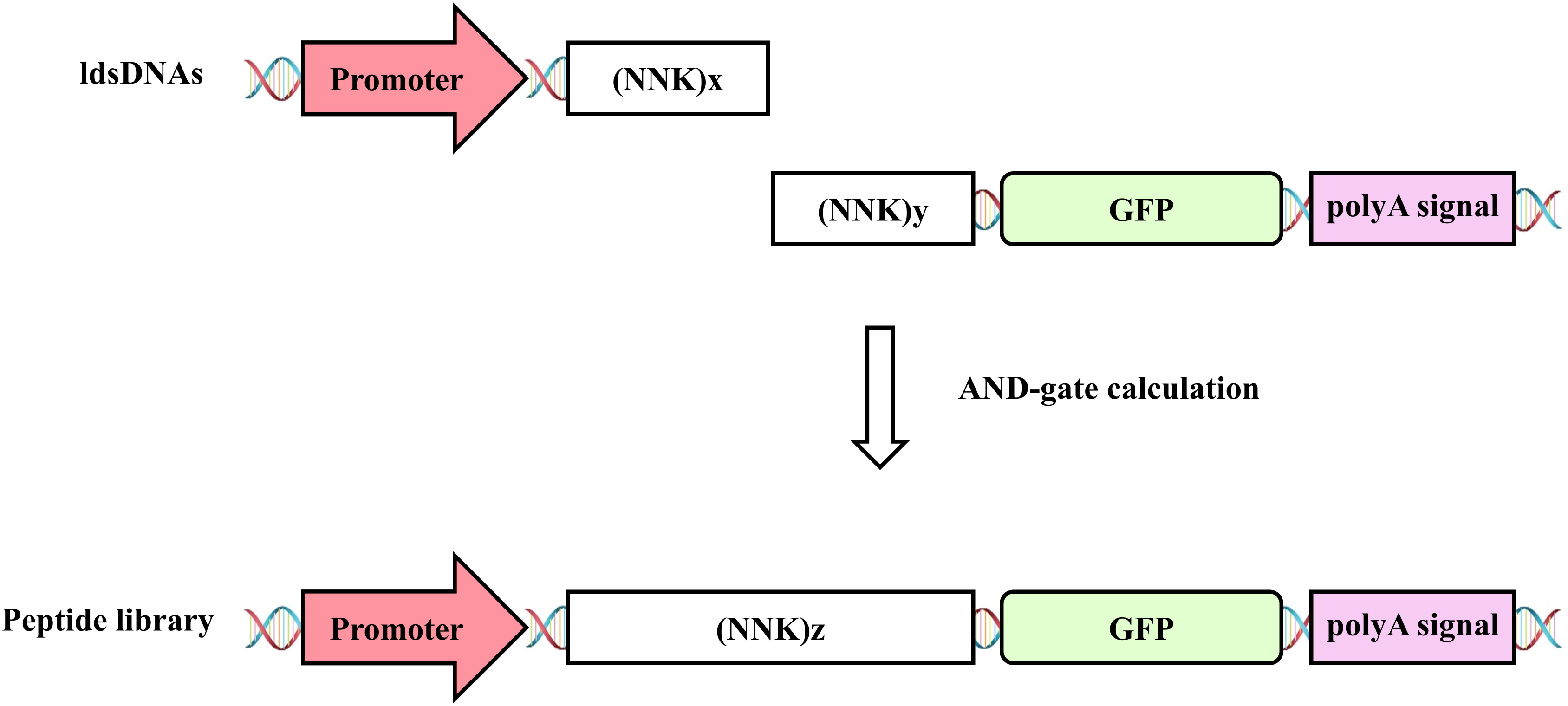
Schematic showing the rationale of generating highly diverse peptide library by ldsDNA-based AND-gate (LBAG) calculation. ldsDNAs containing CMV promoter and tandem NNK degenerate codons are cotransfected with ldsDNAs containing NNK codons, GFP coding region, and polyA signal sequence. The AND-gate calculation is conducted between two kinds of ldsDNA to form peptide library.

Taking pEGFP-C1 plasmid as PCR template, primers with multiple tandem NNK-trinucleotides were used to generate ldsDNAs with terminal degenerate codons (Supplementary Table 1). Three kinds of ldsDNAs, containing CMV promoter, Kozak sequence and different number of NNK-trinucleotides, were produced (namely CMV-Kozak-<NNK>_5_, CMV-Kozak-<NNK>_10_, CMV-Kozak-<NNK>_15_) (for sequences see Supplementary Materials). Meanwhile, two kinds of ldsDNAs, containing terminal NNK-trinucleotides (or not), GFP coding sequence and SV40 polyA signal (PAS), were amplified (namely <NNK>_5_-GFP-SV40 PAS, <NNK>_0_-GFP-SV40 PAS). Six different combinations of these two subgroup ldsDNAs were introduced into HEK293T cells to construct the AND-gate genetic circuits. The six combinations are as following: 1, CMV-Kozak-<NNK>_5_ with <NNK>_5_-GFP-SV40 PAS (5X+5X for short); 2, CMV-Kozak-<NNK>_10_ with <NNK>_5_-GFP-SV40 PAS (10X+5X); 3, CMV-Kozak-<NNK>_15_ with <NNK>_5_-GFP-SV40 PAS (15X+5X); 4, CMV-Kozak-<NNK>_5_ with <NNK>_0_-GFP-SV40 PAS (5X+0X); 5, CMV-Kozak-<NNK>_10_ with <NNK>_0_-GFP-SV40 PAS (10X+0X); 6, CMV-Kozak-<NNK>_15_ with <NNK>_0_-GFP-SV40 PAS (15X+0X).

When AND-gate genetic circuits are built up in cells, peptide library would be generated surrounding the junctions formed between these two subtype ldsDNAs (Figure 1). Forty-eight hours after transfection, total RNAs were extracted then cDNAs were synthesized via reverse transcription. Nucleotide sequences surrounding the junctions formed by two ldsDNA-inputs were PCR amplified and then subjected to pair-end high-throughput sequencing (HTS). The HTS statistics for all sequencing libraries are shown in Table 1.

**Table 1.**
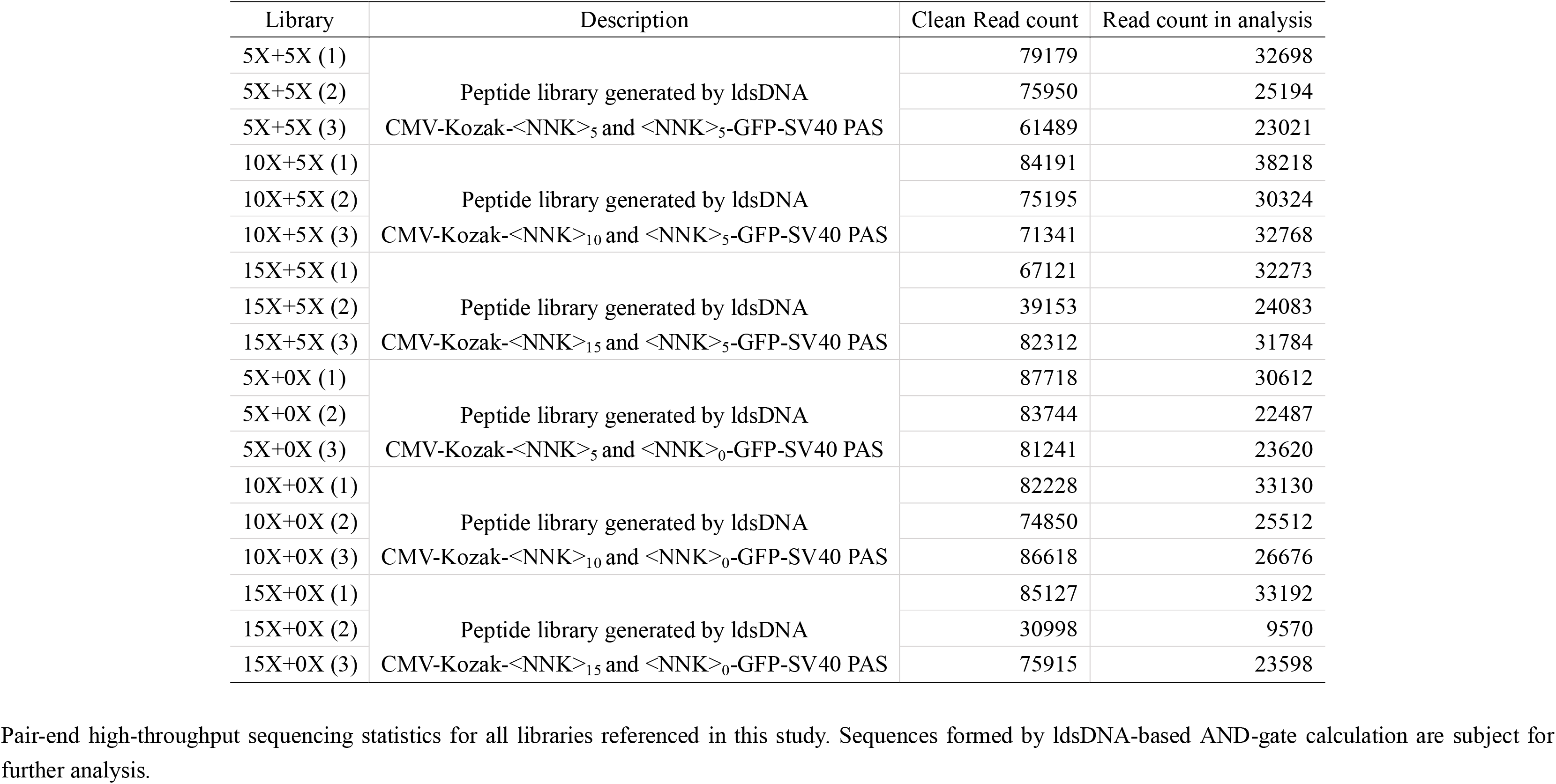
High-throughput sequencing (HTS) statistics

### Reading frame phase of ldsDNA-based peptide library

To display or tether library peptides, oligopeptides are usually fused to specific anchor proteins. For example in phage display experiment, the peptide library is fused to an open reading frame (ORF) thereby connecting peptide to phage coat protein (1). Our previous research demonstrated that the ldsDNA could be subject to terminal nucleotide(s) deletion which may lead to reading frame shift. Here we analyze the reading frame phase characteristics of the LBAG peptide libraries (Figure 2). Libraries generated by dual-side (_X+_X) terminal-NNKs-ldsDNAs, e.g., CMV-Kozak-<NNK>_5_ with <NNK>_5_-GFP-SV40 PAS (5X+5X), show a similar reading frame phase distribution. The average in-frame (3n) ratio ranging from 45% to 47%. While libraries generated by single-side (_X+0X) terminal-NNKs-ldsDNAs, e.g., CMV-Kozak-<NNK>_5_ with <NNK>_0_-GFP-SV40 PAS (5X+0X), show dramatic dynamics of reading frame phase distribution (Figure 2). 5X+0X peptide libraries reach the maximum 69% in-frame rate compared with 63% of 10X+0X and 53% of 15X+0X, respectively. Then, the in-frame nucleotide sequences (3n) are translated into polypeptides for further analysis.

**Figure 2.**
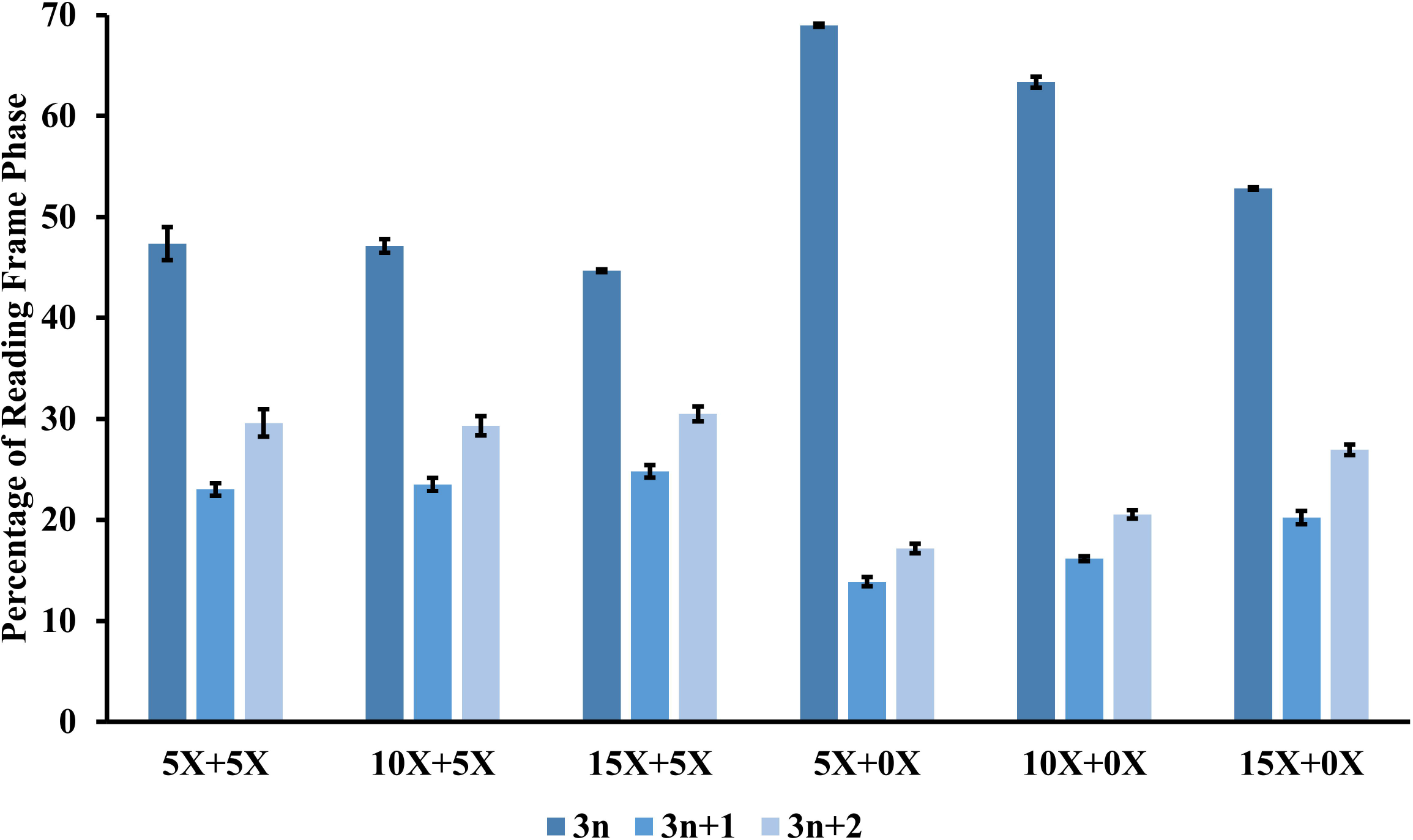
Reading frame character of nucleotide sequences formed by different libraries. Pair-end high-throughput sequencing data is processed to extract the nucleotide sequences formed by ldsDNA-based AND-gate. The nucleotide sequence length is divided by three to show its reading frame character. The average percentage of different phase of different libraries is presented. All data are displayed as the mean ± SD, n = 3.

### Length dynamic and amino acid composition of ldsDNA-based peptide library

We next analyze the polypeptide length distributions of different LBAG peptide libraries (peptides with pre-stop codon are excluded). Dual-side terminal-NNKs-ldsDNAs strategies, including 5X+5X, 10X+5X and 15X+5X, show a more dynamic length distribution compared with that from single-side terminal-NNKs-ldsDNAs strategies (5X+0X, 10X+0X and 15X+0X) (Figure 3). In addition, for single-side terminal-NNKs-ldsDNAs strategies, most peptides have a length equal to the theoretical polypeptide length generated by direct ligation (supposing no insertion or deletion happened) (Figure 3D-F). For example, in the 5X+0X group, over 81% of the peptide length is five amino acids.

**Figure 3.**
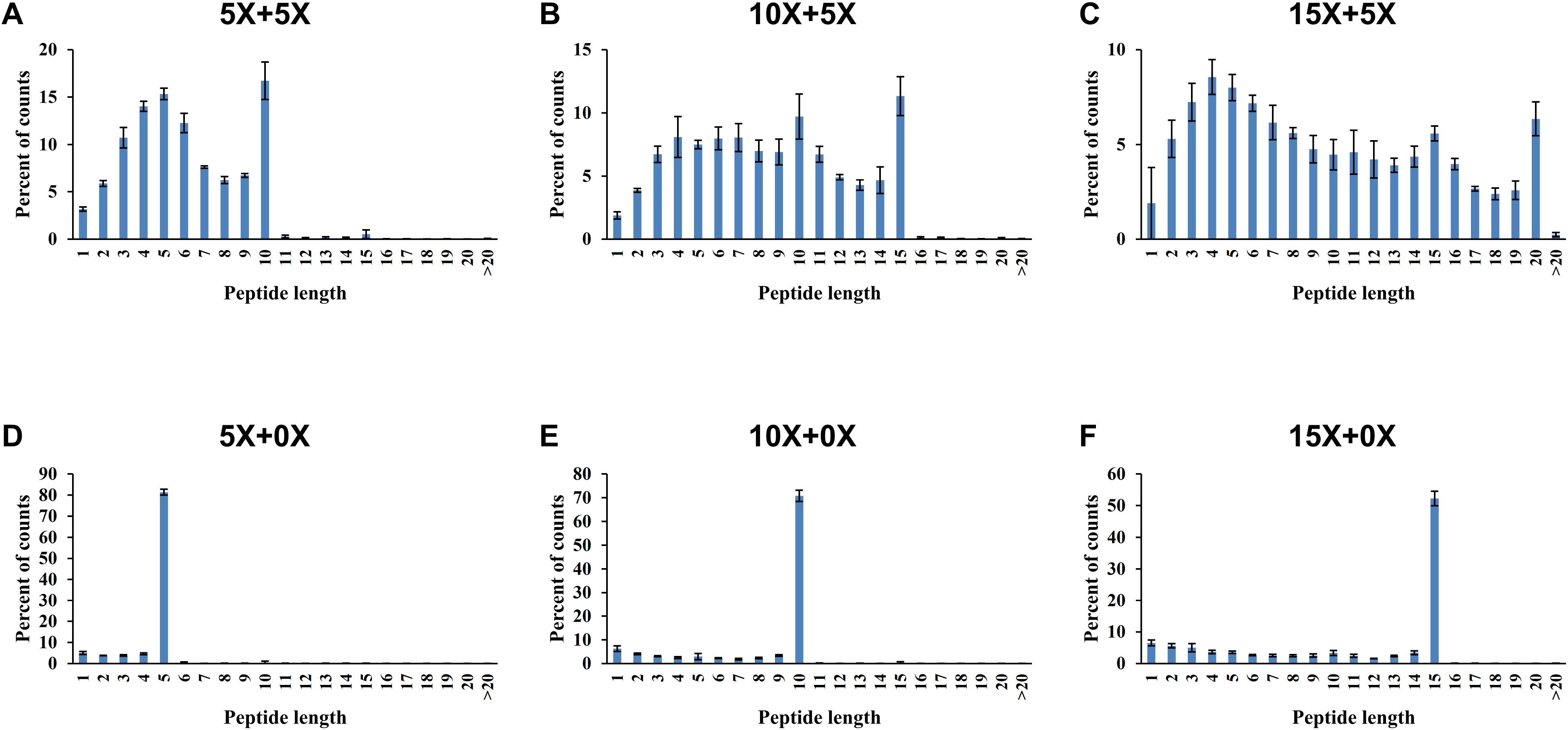
The peptide length distributions of six different peptide libraries generated by ldsDNA-based AND-gate genetic circuits. In-frame (3n) nucleotide sequences are translated into polypeptide sequences. Then the length of polypeptide sequence in each library is summarized. The average percentage of different polypeptide length is presented. All data are displayed as the mean ± SD, n = 3.

The amino acid composition of different LBAG peptide libraries is in line with the theoretical NNK amino acid composition (pre-stop codon is included in the analysis) (Table 2). However, Proline (P) residue always shows a higher percentage than the theoretical ratio, particularly when the peptide chain getting longer. Meanwhile, the percentage of each nucleotide at each position of peptide library codons is showed in Supplementary Table 2. The nucleotide distributions are in line with NNK degenerate base ratio.

**Table 2.** Percentage of each amino acid of different peptide library

| Amino acid | NNK | 5X+5X | 10X+5X | 15X+5X | 5X+0X | 10X+0X | 15X+0X |
| --- | --- | --- | --- | --- | --- | --- | --- |
| Stop codon | 3.125 | 2.34 ± 0.11 | 2.46 ± 0.14 | 2.28 ± 0.15 | 2.59 ± 0.14 | 2.47 ± 0.09 | 2.31 ± 0.13 |
| A (Ala) | 6.25 | 6.16 ± 0.08 | 6.34 ± 0.18 | 7.11 ± 0.19 | 5.43 ± 0.12 | 6.22 ± 0.13 | 7.32 ± 0.26 |
| C (Cys) | 3.125 | 2.80 ± 0.10 | 2.72 ± 0.09 | 2.74 ± 0.24 | 2.06 ± 0.17 | 2.35 ± 0.03 | 2.16 ± 0.09 |
| D (Asp) | 3.125 | 2.84 ± 0.18 | 2.96 ± 0.26 | 2.76 ± 0.07 | 3.72 ± 0.07 | 3.13 ± 0.07 | 2.85 ± 0.05 |
| E (Glu) | 3.125 | 2.74 ± 0.07 | 2.75 ± 0.05 | 2.63 ± 0.11 | 3.67 ± 0.11 | 3.24 ± 0.12 | 3.19 ± 0.10 |
| F (Phe) | 3.125 | 3.34 ± 0.18 | 3.34 ± 0.08 | 2.88 ± 0.08 | 1.71 ± 0.05 | 2.52 ± 0.12 | 1.88 ± 0.04 |
| G (Gly) | 6.25 | 5.23 ± 0.23 | 5.28 ± 0.14 | 5.85 ± 0.08 | 4.35 ± 0.07 | 4.89 ± 0.13 | 5.68 ± 0.14 |
| H (His) | 3.125 | 3.54 ± 0.17 | 3.41 ± 0.18 | 3.26 ± 0.20 | 4.35 ± 0.11 | 3.72 ± 0.14 | 3.21 ± 0.10 |
| I (Ile) | 3.125 | 3.45 ± 0.08 | 3.40 ± 0.38 | 2.76 ± 0.12 | 3.49 ± 0.26 | 3.05 ± 0.12 | 2.48 ± 0.07 |
| K (Lys) | 3.125 | 3.57 ± 0.16 | 3.20 ± 0.06 | 2.70 ± 0.07 | 5.93 ± 0.17 | 4.08 ± 0.09 | 3.32 ± 0.15 |
| L (Leu) | 9.375 | 8.84 ± 0.16 | 9.35 ± 0.17 | 8.98 ± 0.28 | 6.69 ± 0.22 | 8.63 ± 0.11 | 7.93 ± 0.12 |
| M (Met) | 3.125 | 2.91 ± 0.09 | 2.90 ± 0.18 | 2.68 ± 0.03 | 3.54 ± 0.10 | 3.04 ± 0.04 | 2.59 ± 0.08 |
| N (Asn) | 3.125 | 4.23 ± 0.18 | 3.55 ± 0.16 | 2.90 ± 0.07 | 6.33 ± 0.09 | 4.37 ± 0.12 | 3.35 ± 0.12 |
| P (Pro) | 6.25 | 8.16 ± 0.15 | 9.20 ± 0.52 | 10.40 ± 0.28 | 7.34 ± 0.10 | 9.41 ± 0.16 | 11.32 ± 0.11 |
| Q (Gln) | 3.125 | 3.13 ± 0.11 | 3.12 ± 0.04 | 3.17 ± 0.09 | 4.13 ± 0.11 | 3.89 ± 0.02 | 3.82 ± 0.09 |
| R (Arg) | 9.375 | 8.34 ± 0.28 | 8.47 ± 0.24 | 9.79 ± 0.23 | 8.74 ± 0.08 | 9.18 ± 0.25 | 10.68 ± 0.29 |
| S (Ser) | 9.375 | 10.30 ± 0.16 | 9.76 ± 0.30 | 9.68 ± 0.20 | 8.54 ± 0.14 | 8.63 ± 0.17 | 8.90 ± 0.32 |
| T (Thr) | 6.25 | 7.24 ± 0.17 | 6.64 ± 0.27 | 6.94 ± 0.12 | 8.88 ± 0.21 | 7.27 ± 0.17 | 7.58 ± 0.15 |
| V (Val) | 6.25 | 5.66 ± 0.24 | 5.56 ± 0.19 | 5.31 ± 0.21 | 3.59 ± 0.11 | 4.58 ± 0.07 | 4.51 ± 0.21 |
| W (Trp) | 3.125 | 2.30 ± 0.06 | 2.70 ± 0.31 | 2.60 ± 0.04 | 1.96 ± 0.07 | 2.33 ± 0.05 | 2.36 ± 0.07 |
| Y (Tyr) | 3.125 | 2.89 ± 0.11 | 2.89 ± 0.21 | 2.59 ± 0.07 | 2.96 ± 0.11 | 2.99 ± 0.14 | 2.57 ± 0.16 |

In-frame (3n) nucleotide sequences generated by ldsDNA-based AND-gate are translated into polypeptide sequences. The total number of each amino acid in each
library is summarized. The average percentage of different amino acid is presented. All data are displayed as the mean ± SD, n = 3. The theoretical amino acid percentage of NNK degenerate codon is also presented.

### The capacity of the LBAG peptide library

Peptide library capacity could be represented by its coverage ability on amino acid combination possibilities. Here, all eighteen LBAG peptide libraries cover all 400 possible dual-amino-acid combinations (2-mer). For triple-amino-acid (3-mer), the library coverage is ranging from 64% to 98% (Figure 4A). For quadruple-amino-acid (4-mer), the library coverage is ranging from 5.7% to 32.1% (Figure 4B). The peptide library content, defined as the total amino acid number coded by a library, serves as a determinate factor of library capacity (Figure 4).

**Figure 4.**
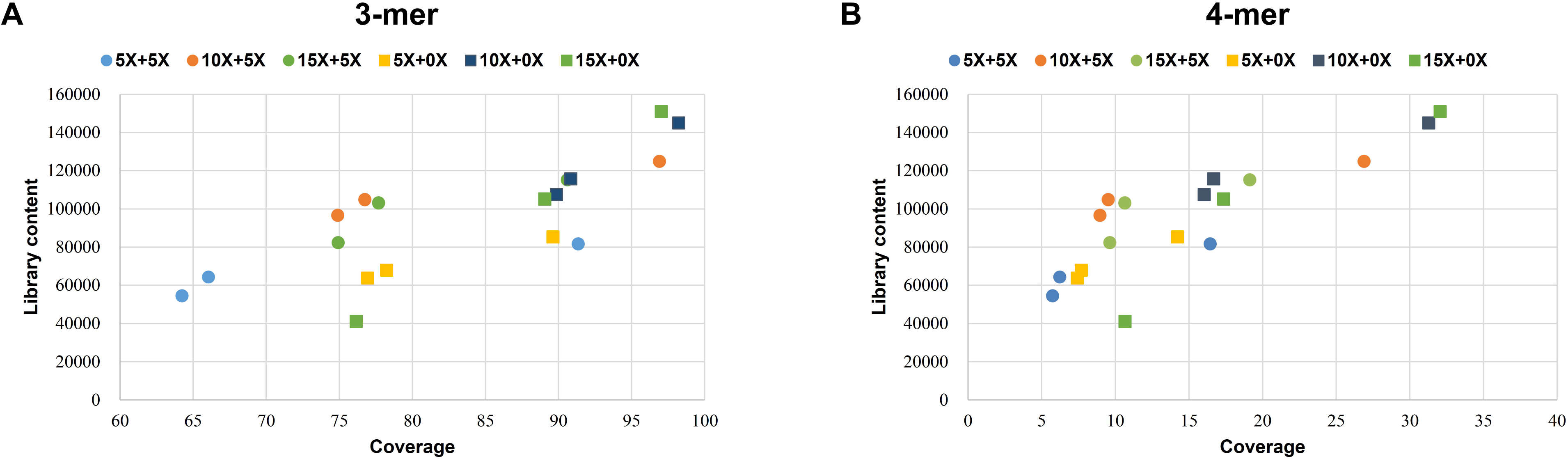
Peptide library coding capacity on covering triple-amino-acid (A) or quadruple-amino-acid (B) peptide possibilities. Peptide libraries are scanned by all triple or quadruple amino-acid combinations, respectively. The library content (defined as the total amino acid number coded by a library) and the percentage of coverage are presented.

### The asymmetry of input ldsDNA end processing during AND gate calculation

As shown by Figure 2 and 3, single-side (_X+0X) and dual-side (_X+_X) LBAG peptide library strategies show obvious differences in both reading frame phase distributions and peptide length dynamics. To further investigate these differences and figure out whether or not the up-stream and down-stream ldsDNAs have the same chance of end processing, we PCR-generated two pairs of AND gate ldsDNAs: pair 1, CMV-Kozak-FLAG-NNN with GFP-SV40 PAS; and pair 2, CMV-Kozak-FLAG with NNN-GFP-SV40 PAS (N for randomized nucleotides, for ldsDNA sequences see Supplementary Materials). After cell transfection, RNA extraction, cDNA synthesis, and PCR amplification, high-throughput sequencing (HTS) was performed to characterize the oligonucleotide sequences formed by AND gate calculation. About 67% of the sequences formed by CMV-Kozak-FLAG-NNN and GFP-SV40 PAS contains intact random nucleotides (NNN) (Figure 5). While for CMV-Kozak-FLAG and NNN-GFP-SV40 PAS group, about 57% of oligonucleotide contains triple nucleotides (Figure 5). Thus, the down-stream ldsDNAs (NNN-GFP-SV40 PAS) are more likely subject to end processing than up-stream ldsDNAs (CMV-Kozak-FLAG-NNN) in AND gate relink process in cells. This may be the reason why single-side (_X+0X) strategy generate higher in-frame rate and less variability among the peptide length than dual-side (_X+_X) strategy.

**Figure 5.**
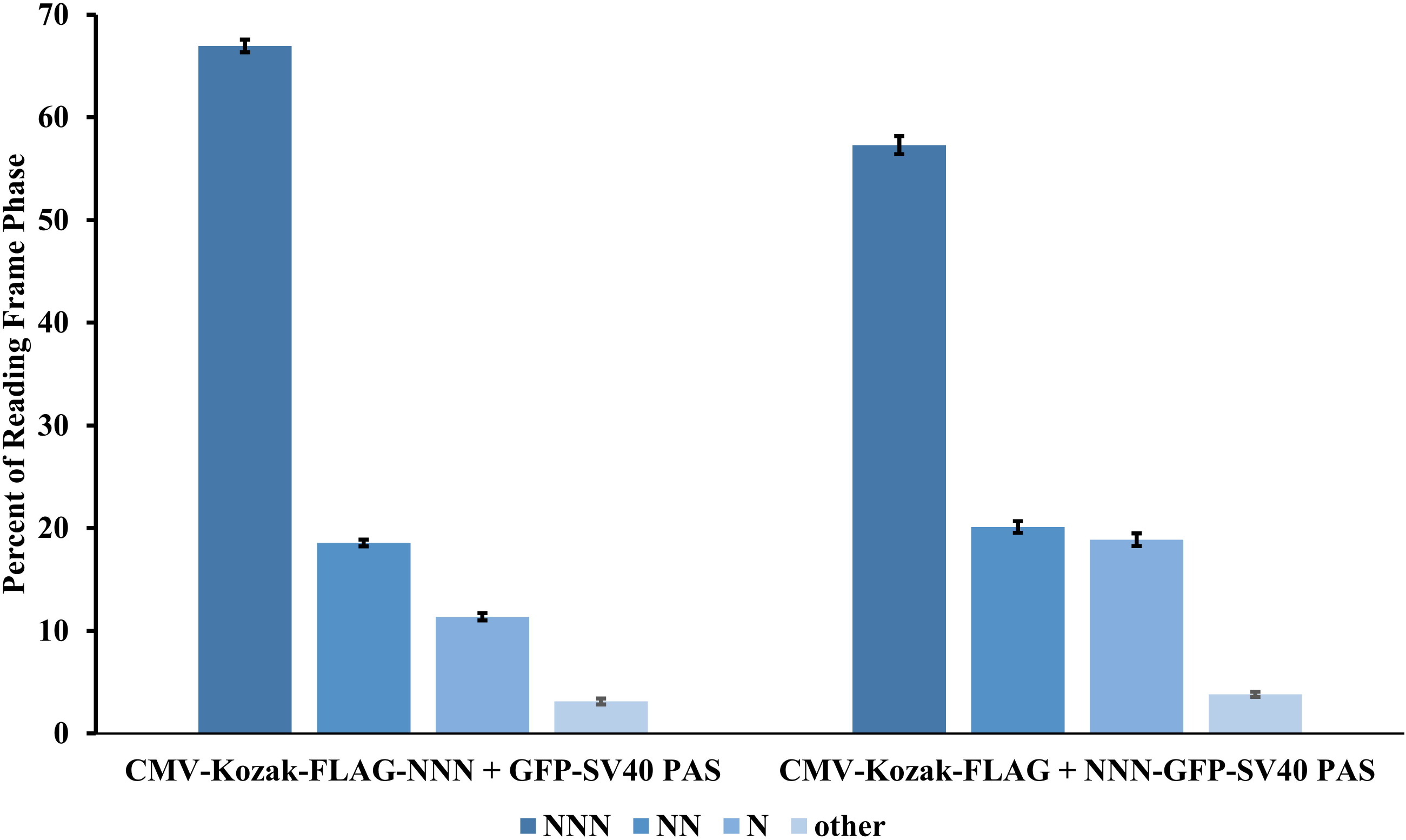
Reading frame character of nucleotide sequences formed by ldsDNAs with terminal-triple-random-nucleotides. Two pairs of AND gate ldsDNAs were PCR-generated and transfected into HEK293T cells (pair 1, CMV-Kozak-FLAG-NNN with GFP-SV40 PAS; pair 2, CMV-Kozak-FLAG with NNN-GFP-SV40 PAS). Pair-end high-throughput sequencing data is processed to extract the nucleotide sequences formed by ldsDNA-based AND-gate. The average percentage of different phase of different libraries is presented. All data are displayed as the mean ± SD, n = 4.

In addition, through the above terminal-triple-random-nucleotide-ldsDNA AND gate calculation, we also evaluated the nucleotide preferences of LBAG calculation. In both CMV-Kozak-FLAG-NNN + GFP-SV40 PAS group and CMV-Kozak-FLAG + NNN-GFP-SV40 PAS group, Guanine (G) is always the unfavorable end nucleotide (Supplementary Table 3). Furthermore, up-stream or down-stream ldsDNAs containing two or three terminal Guanines are hard to connect to the corresponding ldsDNAs (Supplementary Table 3).

### Using LBAG peptide library to identify MDM2 protein interacting peptide

The E3 ubiquitin ligase MDM2 negatively regulates the activity of the tumor suppressor protein p53 through interacting with the N-terminal transactivation domain of p53. Structurally, the residues F19, W23 and L26 in p53, which insert deep into the MDM2 hydrophobic cleft, are critical in MDM2-p53 interaction (21). Also, the non-peptide inhibitors of the p53-MDM2 interaction have to contain mimics of these amino acids (22–24). Pazgier *et al.* screened a duodecimal peptide phage display library to identify MDM2 (residues 25-109) interacting peptides (25). They found 14 out of 15 MDM2 binding clones containing the same conserved residues: <u>**F**</u>XXX<u>**W**</u>XX<u>**L**</u>, which in line with the MDM2-p53 binding pattern. Shiheido *et al.* employed mRNA display to identify MDM2 interacting peptides (26,27). Similarly, they found more than half of all peptides retained the three hydrophobic residues corresponding to F19, W23 and L26 of wild-type p53. Here, to test the performance of our LBAG peptide library strategy, we plan to identify the MDM2 interacting peptides in mammalian cells.

To detect protein-protein interaction in mammalian cells, we employed the two-hybrid system, which has been adapted for use in mammalian cells (28,29). The system contains three plasmids: pACT, which expresses VP16 activation domain, pBIND, which expresses GAL4 DNA-binding domain, pG5egfp, which contains five GAL4 binding sites upstream of a minimal TATA box and EGFP ORF. In the present study, MDM2 (residues 25-109) is subcloned into pACT plasmid (namely pACT-MDM2). By PCR amplification, the LBAG peptide library is fused to GAL4 DNA-binding domain through an XTEN linker (Figure 6A). To achieve high library content, we here used single-side LBAG strategy.

**Figure 6.**
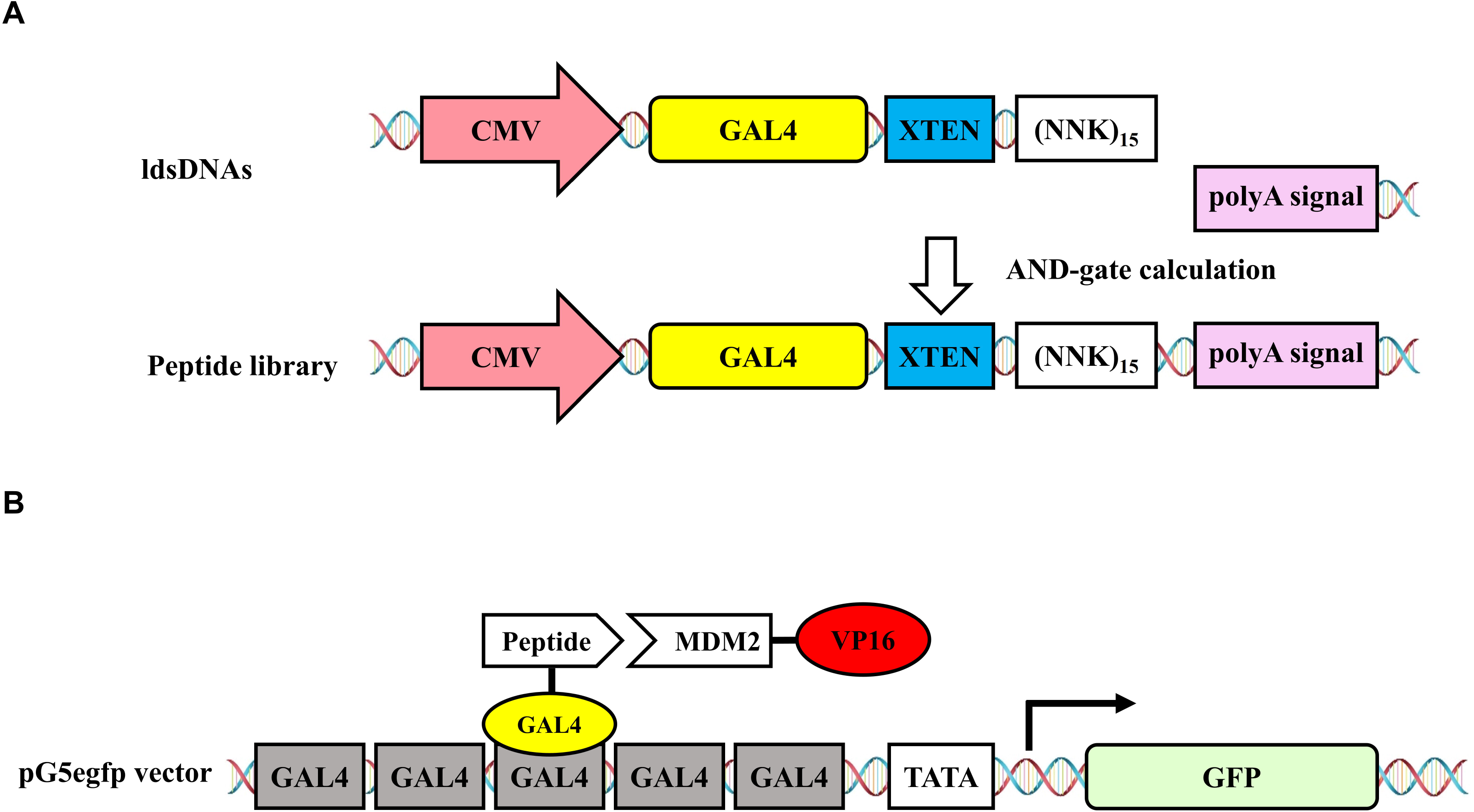
Schematic showing the rationale of generating LBAG peptide library for two-hybrid system in mammalian cells. (A) ldsDNA CMV-GAL4-XTEN-<NNK>_15_ (containing CMV promoter, GAL4-XTEN coding sequence, tandem NNK degenerate codons) is cotransfected with ldsDNA SV40 PAS. The AND-gate calculation is conducted between two ldsDNAs to form peptide library. (B) Schematic representation of the mammalian two-hybrid system. The pG5egfp plasmid contains five GAL4 binding sites upstream of a minimal TATA box, which in turn is upstream of the GFP gene. Interaction between the two test proteins, GAL4-peptide and MDM2-VP16, results in an expression of GFP protein.

Two plasmids (pACT-MDM2, pG5egfp) and two ldsDNAs (CMV-GAL4-XTEN-<NNK>_15_, SV40 PAS) were cotransfected into HEK293T cells. The peptide library will be formed through the AND gate calculation between CMV-GAL4-XTEN-<NNK>_15_ and SV40 PAS (Figure 6A). If the peptide interacts with MDM2, the activation domain VP16 will be recruited to GAL4 binding sites, thereby activating the GFP reporter gene expression (Figure 6B). Here fluorescence-activated cell sorting (FACS) was used to isolate the GFP positive cells. High-throughput sequencing of the peptide library in these cells identified peptide S<u>**F**</u>VIR<u>**W**</u>TA<u>**L**</u>WPGPRS among the top hits (23 reads, rank fourth) (Supplementary Table 4). This peptide contains the conserved residues <u>**F**</u>XXX<u>**W**</u>XX<u>**L**</u>, which is critical to interact with MDM2.

## Discussion

Affinity-based display technologies, e.g., phage, ribosome, mRNA, bacterial, and yeast-display, have been developed to screen peptide libraries to identify protein-binding peptides. Peptide library size is a determinate factor of a success screen. For phage and yeast-display, the complexity of peptide library stores in plasmid (1,2,7). Currently, the phage display library size could reach 10^9^ (9,30). While for ribosome and mRNA-display, PCR products serve as genetic medium to store library coding information. Ribosome and mRNA-display could achieve extremely large library size (>10^13^) (5,31). In the current research, we provide a new strategy to build high content peptide library in cultured cell line. The protocol is easy to conduct, only containing two steps: PCR amplification and cell transfection. Since degenerate trinucleotides is present at the 5’ end of the primer, the library complexity keeps on increasing during PCR amplification process. Theoretically, no specific nucleotide sequence is enriched during PCR reaction. PCR products are then transfected into cells to conduct AND-gate linkage to generate full gene expression cassette (promoter-CDS-PAS). For our strategy, no particular attention needs to be paid to maintain the library abundance. The size of ldsDNA-based library needs to be further investigated. Determining the peptide sequence diversity in each cell by single-cell-sequencing will provide crucial information for estimating the whole library size.

For the current protocol, there are still several issues that need to be further addressed to increase library capacity. First of all, the high in-frame ratio would improve library capacity. Our results reveal that peptide libraries generated by single-side NNKs-ldsDNA show higher in-frame ratios than that from dual-side NNKs-ldsDNAs (Figure 2). Besides, chemical modified primers could be used to generate nuclease resistance ldsDNAs to prevent reading frame shift (32,33). Another major obstacle is the pre-stop codon, which causes premature termination (34). For NNK degenerate codon, the percentage of stop-codon in each amino acid position is 3.125% (1/32) and will be accumulated when the peptide is getting longer. When the polypeptide length reaches 20 amino acids, about 47% of the sequences contain pre-stop codon. One potential solution to avoid stop codon is to synthesize PCR primer using trimer phosphoramidites (35,36).

In the present study, we tested the feasibility of our LBAG peptide library strategy through the identification of the MDM2 interacting peptide (Figure 6). Indeed, one peptide (S<u>**F**</u>VIR<u>**W**</u>TA<u>**L**</u>WPGPRS), which contains three crucial residues mediating MDM2-p53 interaction, is among the top hits (Supplementary Table 4). However, in phage display and mRNA display experiments, multiple peptides with <u>**F**</u>XXX<u>**W**</u>XX<u>**L**</u> pattern were identified (25–27). This may come from the enrichment of interacting peptides during rounds of panning in phage and mRNA display process. For the current protocol, four components, including two plasmids (pACT-MDM2, pG5egfp) and two ldsDNAs (CMV-GAL4-XTEN-<NNK>_15_, SV40 PAS), were transient cotransfected into one HEK293T cell to conduct AND gate calculation and assemble the two-hybrid system. In the future application, to save the cell transfection capacity, the plasmids should be preset into cells by generating a stable cell line. Thereby, only ldsDNAs need to be transfected into cells.

Here by terminal-NNKs-<u>**l**</u>dsDNA-<u>**b**</u>ased <u>**A**</u>ND-<u>**g**</u>ate (LBAG) genetic circuit, we develop a novel method to generate highly diverse peptide library in mammalian cells. Our protocol is easy to conduct only PCR-reaction and cell-transfection are needed. High-throughput sequencing results reveal that LBAG peptide library is characterized by both amino acids sequence diversity and peptide length dynamics. We also identified a peptide with conserved residues which are critical to bind to MDM2. Our new method may have great application potential in therapeutic peptide development.

## Materials and Methods

### Cell culture

The HEK293T cell line was maintained in Dulbecco’s modified Eagle’s medium (DMEM) (Thermo Fisher Scientific, Hudson, NH, USA) containing 10% fetal bovine serum (FBS) with 1% penicillin-streptomycin solution at 37 degree with 5% CO_2_.

### ldsDNA synthesis

KOD-Plus-Ver.2 DNA polymerase (Toyobo, Osaka, Japan) was used to amplify ldsDNAs (PCR amplicons) using the pEGFP-C1 (Clontech, Mountain View, CA, USA) plasmid as the template. PCR products underwent agarose electrophoresis and gel-purification to remove plasmid template and free dNTPs (Universal DNA purification kit, Tiangen, Beijing, China). The amount of PCR products was determined by the OD 260 absorption value. PCR primer sequences and ldsDNA sequences were present in Supplementary Table 1 and Supplementary Materials, respectively.

### ldsDNA transfection

HEK293T cells were seeded into 6-well plates the day before transfection (60% − 70% confluency). 500 ng/well of each input amplicons (total 1000 ng/well) were transfected into cells with Lipofectamine 2000 (Invitrogen, Carlsbad, CA, USA) reagent following standard protocol.

### RNA extraction, cDNA synthesis and PCR amplification

Forty-eight hours after transfection, total RNAs were extracted by RNAiso Plus (TaKaRa, Beijing, China) reagent. cDNAs were synthesized using the PrimeScript™ RT reagent Kit with gDNA Eraser (TaKaRa) following standard protocol. KOD-Plus-Ver.2 DNA polymerase (Toyobo) was used to amplify the corresponding cDNA sequences surrounding ldsDNA junctions.

### Two-hybrid system in mammalian cells

The two-hybrid system used in this study was developed from commercial CheckMate™ Mammalian Two-Hybrid System (Promega, Madison, WI, USA) with some modifications. Briefly, MDM2 (residues 25-109) was subcloned into pACT (pACT empty vector, GenBank Accession Number AF264723) vector through XbaI and KpnI restriction sites (namely pACT-MDM2). EGFP sequence was cloned into pG5luc vector (pG5luc empty vector, GenBank Accession Number AF264724) to replace luciferase sequence through HindIII and FseI restriction sites (namely pG5egfp). Two ldsDNAs CMV-GAL4-XTEN-<NNK>_15_ and SV40 PAS were PCR-generated using the pBIND (pBIND empty vector, GenBank Accession Number AF264722) plasmid as the template. pACT-MDM2 (500ng/well), pG5egfp (500ng/well), ldsDNA CMV-GAL4-XTEN-<NNK>_15_ (500ng/well) and ldsDNA SV40 PAS (250ng/well) were transient cotransfected into 6-well plated HEK293T cells using Lipofectamine 2000. The experiment could be scaled up by transfected more 6-well plated cells. Forty-eight hours after transfection, FACS was used to isolate the GFP positive cells.

### Library construction and high-throughput sequencing

Library construction and high-throughput sequencing were conduct by Novogene (Beijing, China). Sequencing libraries were generated using TruSeq® DNA PCR-Free Sample Preparation Kit (Illumina, San Diego, CA, USA) following the manufacturer’s recommendations and index codes were added. The library quality was assessed on the Qubit Fluorometer and Agilent Bioanalyzer 2100 system. At last, the libraries were sequenced on an Illumina HiSeq 2500 platform and 250 bp paired-end reads were generated.

### Data analysis

Paired-end reads from the original DNA fragments were merged using FLASH, a very fast and accurate analysis tool which was designed to merge paired-end reads when there are overlaps between reads 1 and reads 2. Paired-end reads were assigned to each sample according to the unique barcodes. Reads were filtered by QIIME quality filters. Nucleotide sequences generated by ldsDNA-based AND-gate genetic circuits were extracted by in-house Perl scripts. High-throughput sequencing data are deposited on the Gene Expression Omnibus (GEO, http://www.ncbi.nlm.nih.gov/geo) under the accession number GEO: GSE134671 (reviewer access token: kpujmequhnspngx).

## Acknowledgements

This work was supported by the National Natural Science Foundation of China [31870860, 31400673 to S.L., 31971388, 81402407 to W.S.]; the Tianjin Research Program of Application Foundation and Advanced Technology [14JCQNJC09800 to S.L., 15JCQNJC11700 to W.S.]; the State Key Laboratory of Medicinal Chemical Biology (Nankai University) [2018103 to S.L.]; and the Key Project of National Health and Family Planning Commission of Tianjin [2017057 to C.Z.]. The authors have declared that no competing interest exists.

## Author Contributions

S.L. conceived and supervised the study. S.L. and W.S. designed and performed the experiments. S.L. and W.S. analyzed the data. W.S. and S.L. prepared the figures. S.L. and W.S. wrote the paper. All authors discussed the results and reviewed the manuscript.

---

**Supplementary Table 1.**
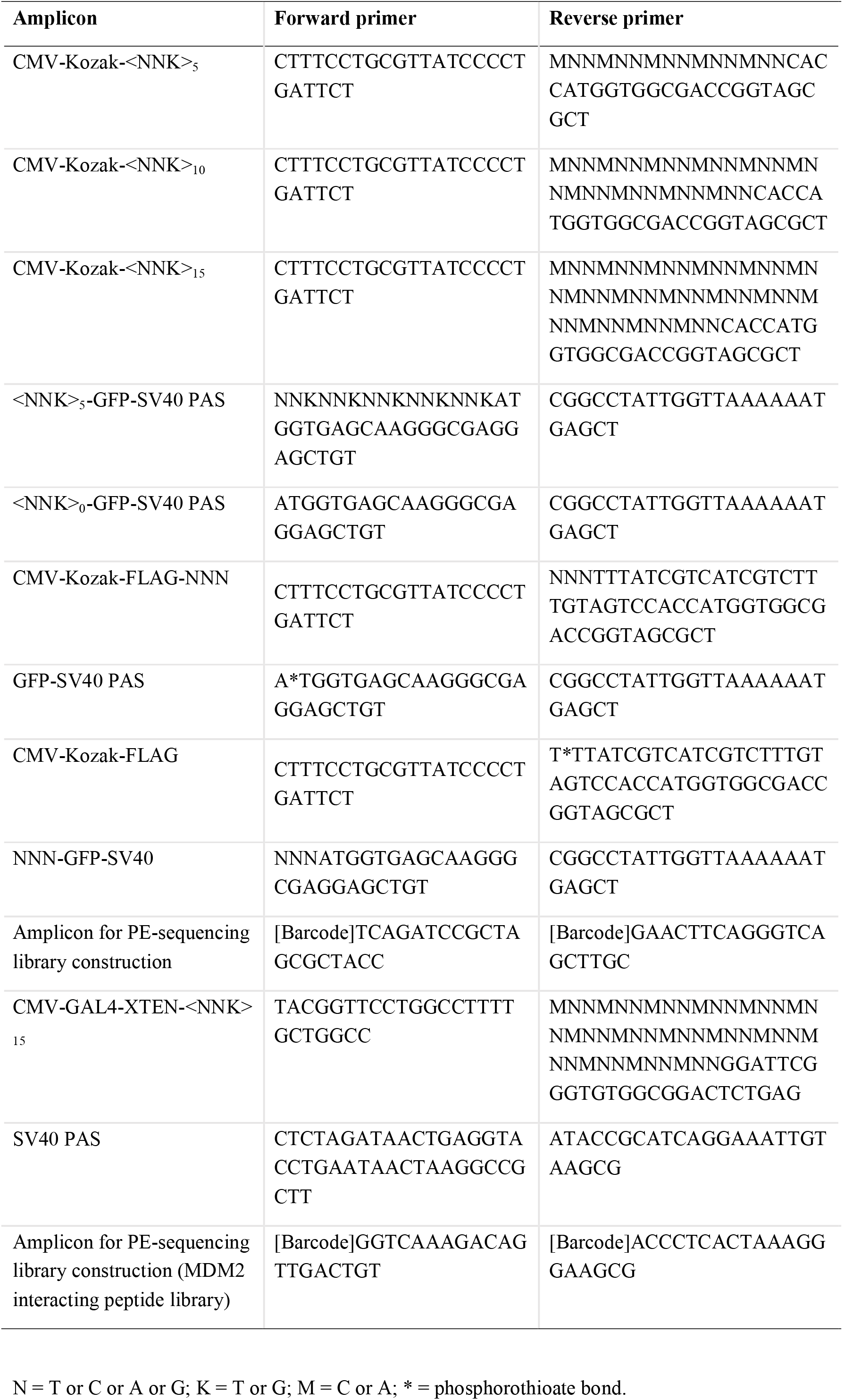
List of PCR primers used in study.

**Supplementary Table 2.** Percentage of each nucleotide at each position of peptide library codons

| Codon position | Nucleotide | 5X+5X | 10X+5X | 15X+5X | 5X+0X | 10X+0X | 15X+0X |
| --- | --- | --- | --- | --- | --- | --- | --- |
| First | T | 23.69 ± 0.30 | 24.31 ± 0.45 | 22.86 ± 0.35 | 17.96 ± 0.45 | 21.21 ± 0.21 | 19.67 ± 0.36 |
|  | C | 26.43 ± 0.30 | 28.23 ± 0.59 | 30.39 ± 0.29 | 25.64 ± 0.34 | 29.42 ± 0.28 | 31.93 ± 0.41 |
|  | A | 27.26 ± 0.23 | 24.58 ± 0.35 | 23.09 ± 0.13 | 35.64 ± 0.22 | 27.31 ± 0.27 | 24.84 ± 0.15 |
|  | G | 22.63 ± 0.18 | 22.89 ± 0.30 | 23.66 ± 0.24 | 20.75 ± 0.11 | 22.06 ± 0.24 | 23.55 ± 0.42 |
| Second | T | 24.20 ± 0.40 | 24.55 ± 0.67 | 22.60 ± 0.10 | 19.01 ± 0.02 | 21.82 ± 0.24 | 19.39 ± 0.35 |
|  | C | 28.78 ± 0.19 | 29.43 ± 0.80 | 31.66 ± 0.35 | 26.53 ± 0.31 | 28.91 ± 0.45 | 32.50 ± 0.29 |
|  | A | 25.23 ± 0.38 | 24.29 ± 0.39 | 22.19 ± 0.34 | 33.66 ± 0.07 | 27.87 ± 0.39 | 24.55 ± 0.50 |
|  | G | 21.79 ± 0.29 | 21.74 ± 0.10 | 23.55 ± 0.09 | 20.80 ± 0.35 | 21.40 ± 0.33 | 23.55 ± 0.13 |
| Third | T | 52.50 ± 0.42 | 51.13 ± 0.24 | 47.80 ± 0.04 | 49.64 ± 0.74 | 48.56 ± 0.34 | 45.07 ± 0.44 |
|  | C | 0.65 ± 0.04 | 0.66 ± 0.06 | 1.51 ± 0.03 | 0.48 ± 0.05 | 0.31 ± 0.03 | 0.83 ± 0.08 |
|  | A | 0.86 ± 0.10 | 0.61 ± 0.04 | 1.03 ± 0.08 | 0.42 ± 0.03 | 0.36 ± 0.04 | 0.60 ± 0.05 |
|  | G | 45.98 ± 0.31 | 47.60 ± 0.29 | 49.66 ± 0.12 | 49.46 ± 0.71 | 50.77 ± 0.33 | 53.50 ± 0.33 |

The total number of each nucleotide at each position of peptide library codons is summarized. The average percentage of different nucleotide at each codon base is presented. All data are displayed as the mean ± SD, n = 3.

**Supplementary Table 3.**
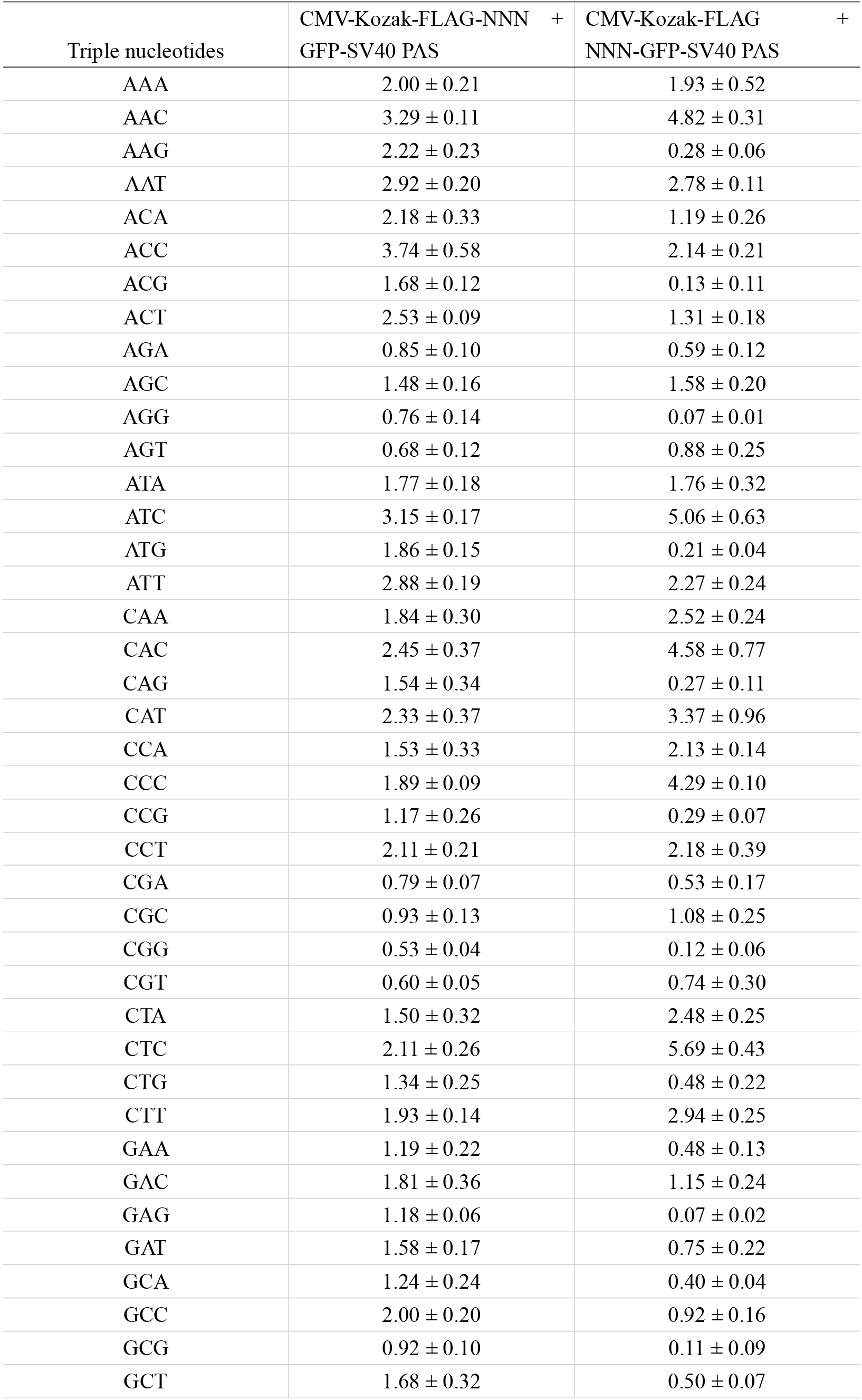
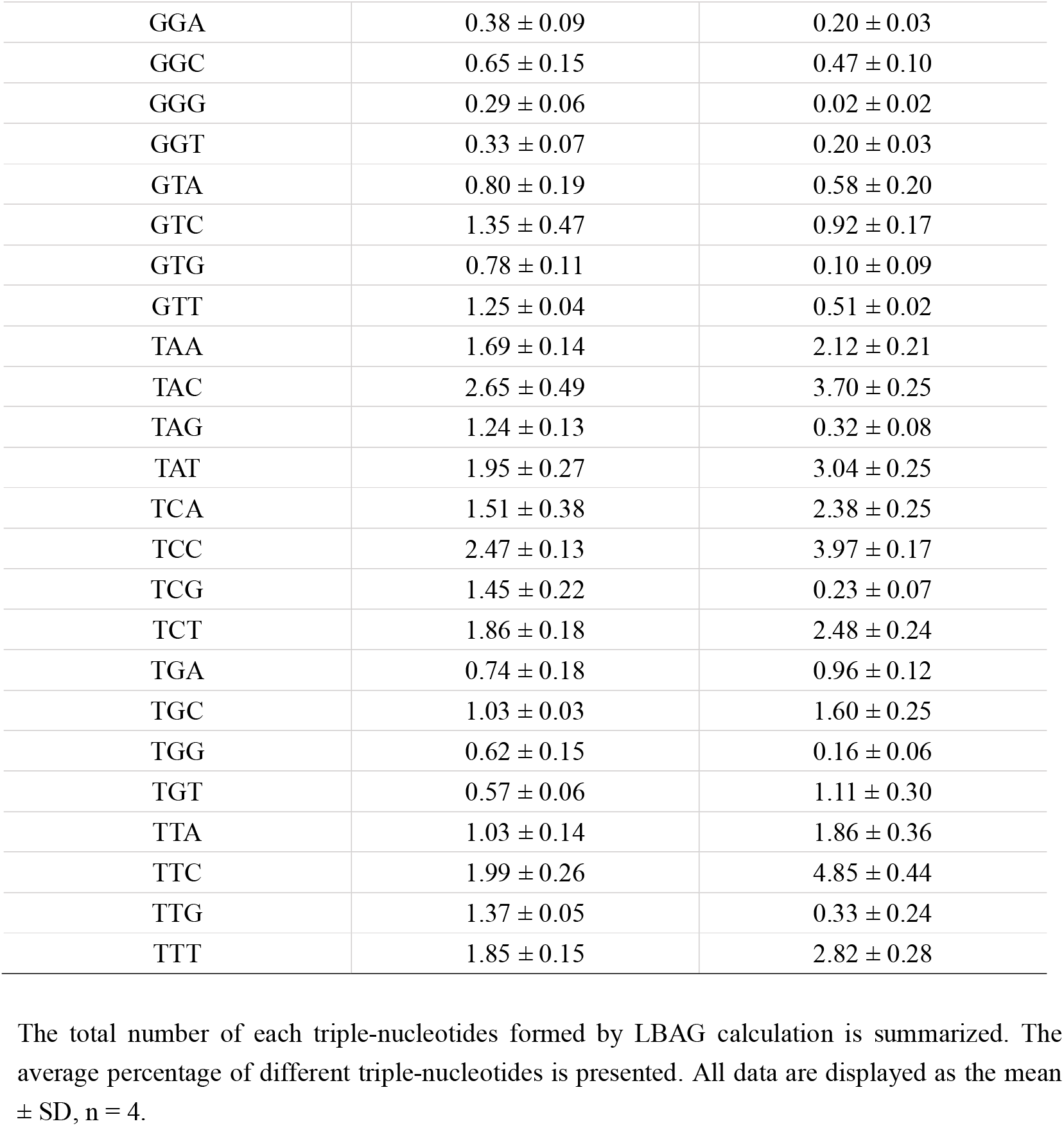
Percentage of triple-nucleotides formed by LBAG calculation

**Supplementary Table 4.** Potential MDM2 interacting peptide

| No | Peptide sequence | Read counts |
| --- | --- | --- |
| 1 | C A G M I K Q S Q L R H P K S | 47 |
| 2 | K L R I M M Q A F R H V Y W N | 39 |
| 3 | D R Q T N Y G N Q L P A R D V | 25 |
| 4 | S <u>F</u> V I R <u>W</u> T A <u>L</u> W P G P R S | 23 |
| 5 | M Y S K W D P M R H D F S E T | 21 |
| 6 | H S L D L Q N Q I Q T C E E W | 20 |
| 7 | S T P W L A M M L R L G P F N | 17 |
| 8 | N S C R L M L H P Q P M R P L | 16 |
| 9 | P T S R E F C N A I H A C L V | 14 |
| 10 | K Y F C M P E T R M H P R R T | 14 |

High-throughput sequencing decoded top ten hit amino acid sequences in GFP positive cells obtained from the LBAG mammalian two-hybrid system. Three crucial residues (Phe, Trp and Leu) mediating MDM2-p53 interaction are shown in underline bold.

## Supplementary materials

### ldsDNA sequences in study

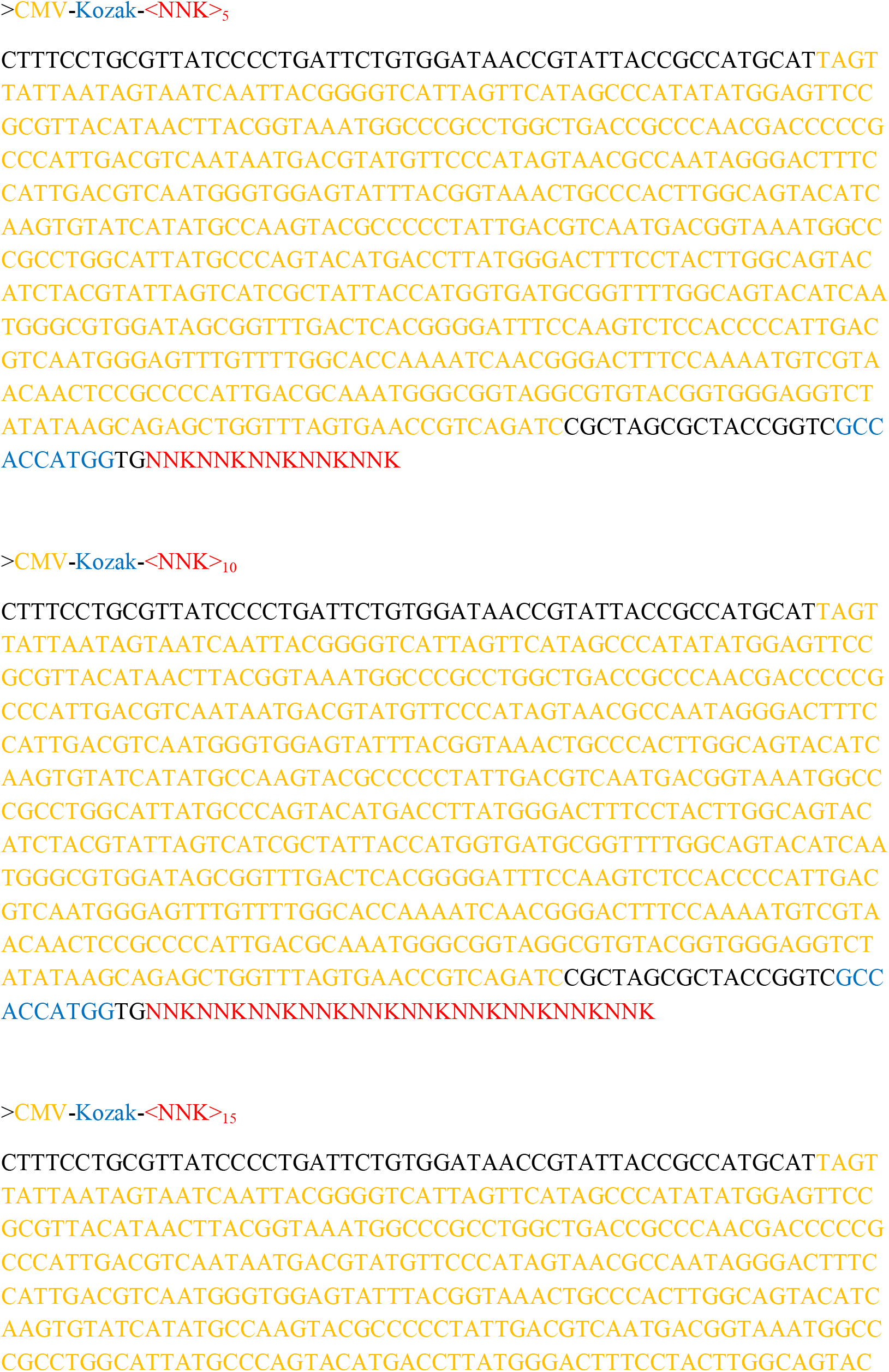

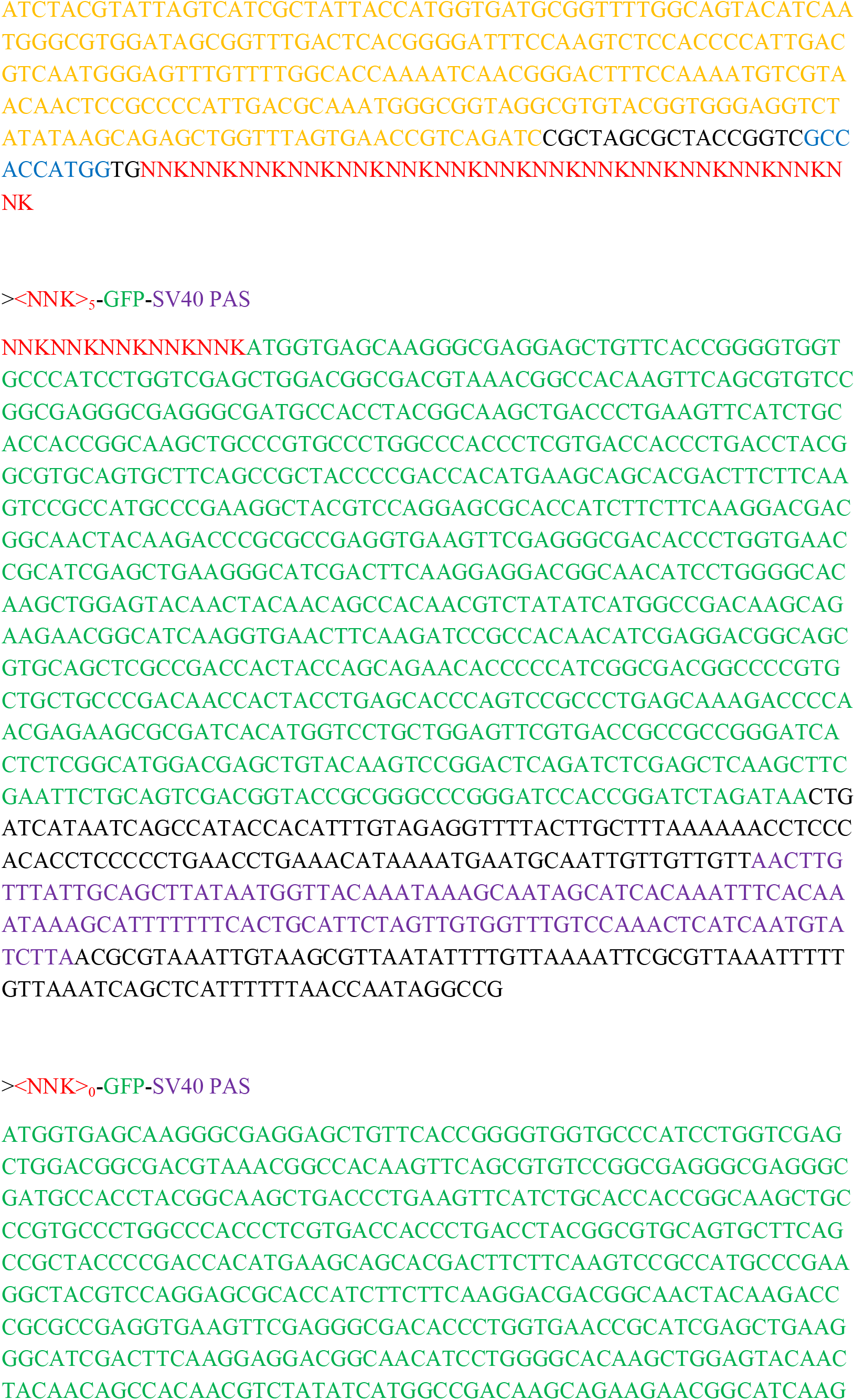

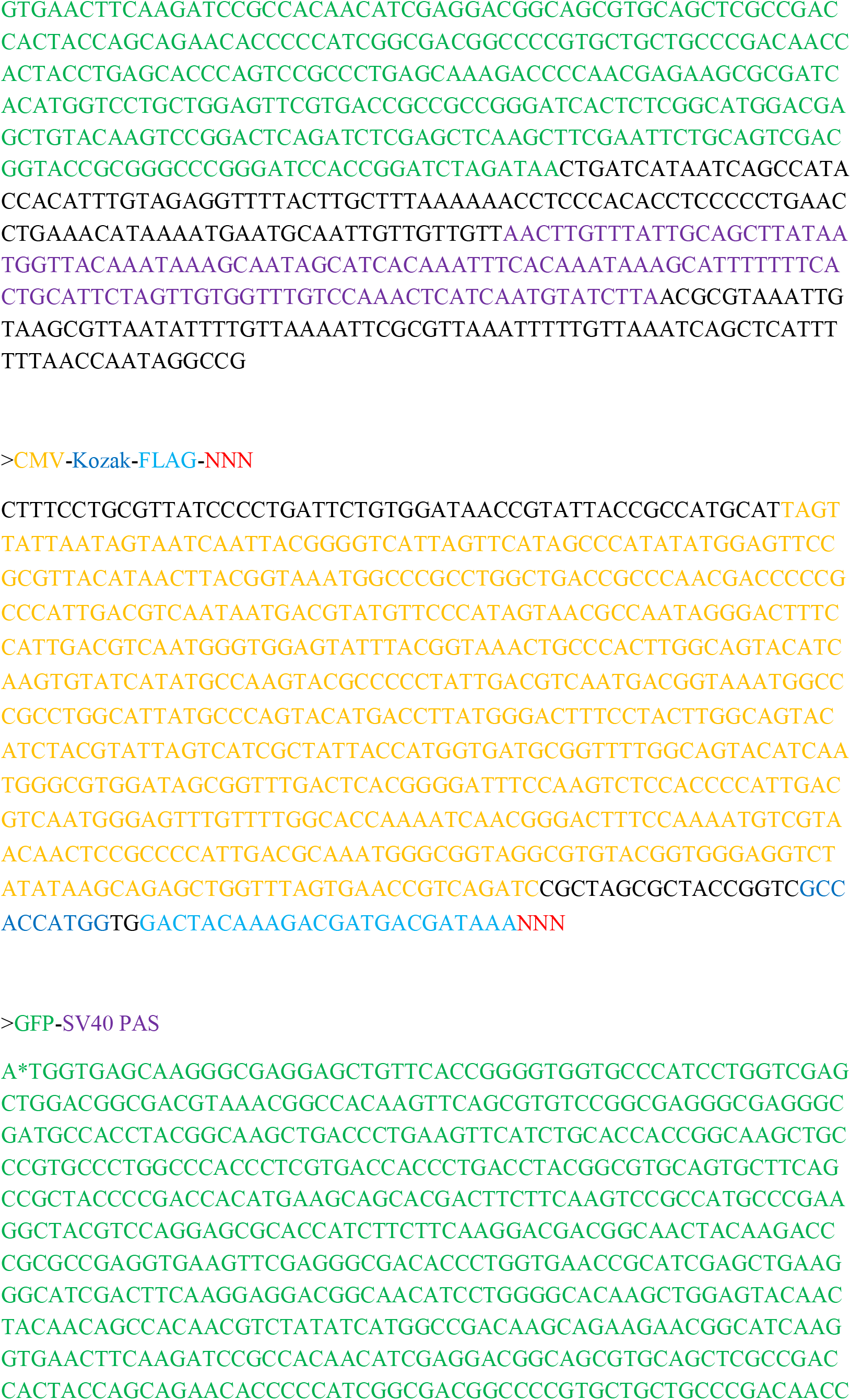

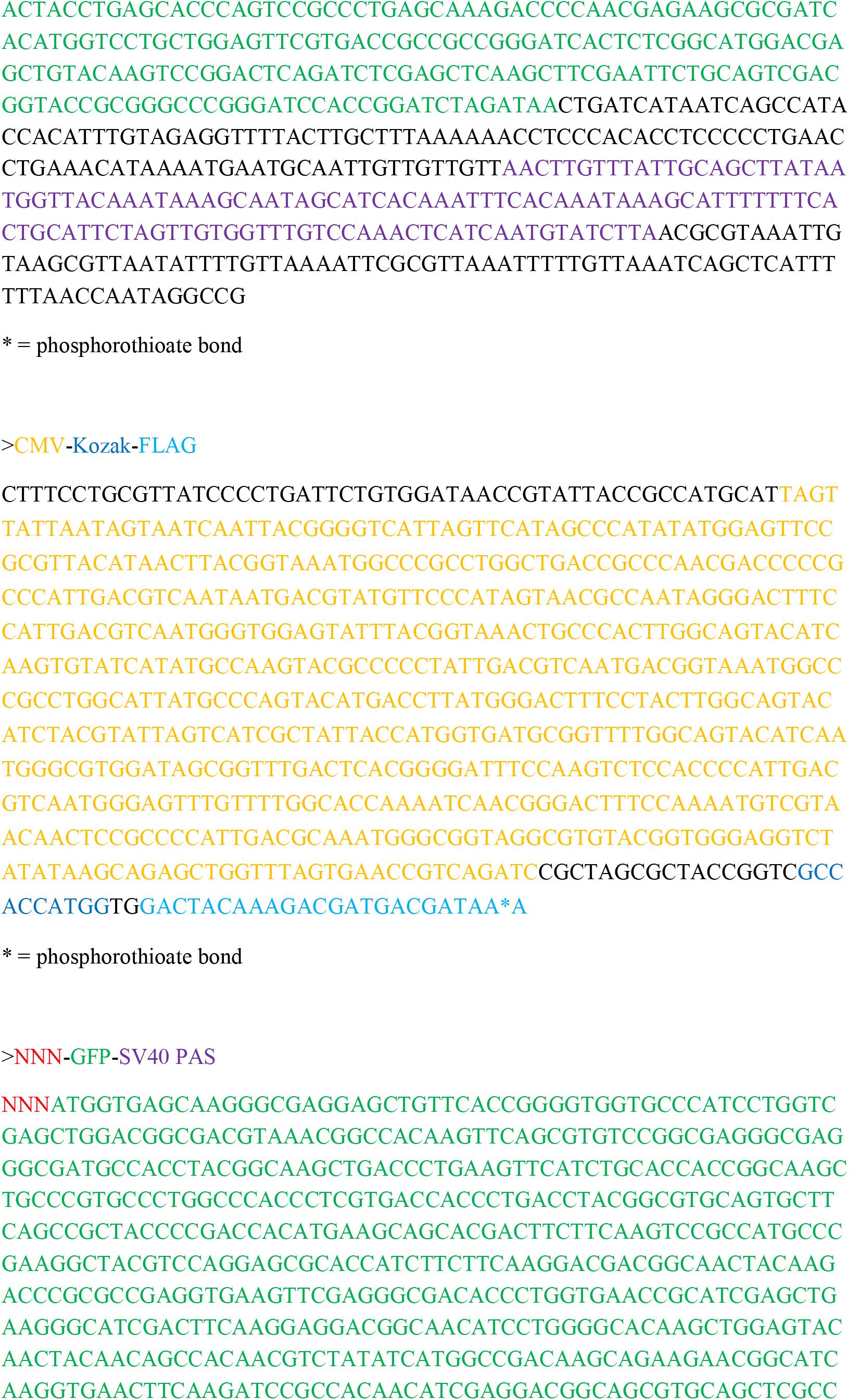

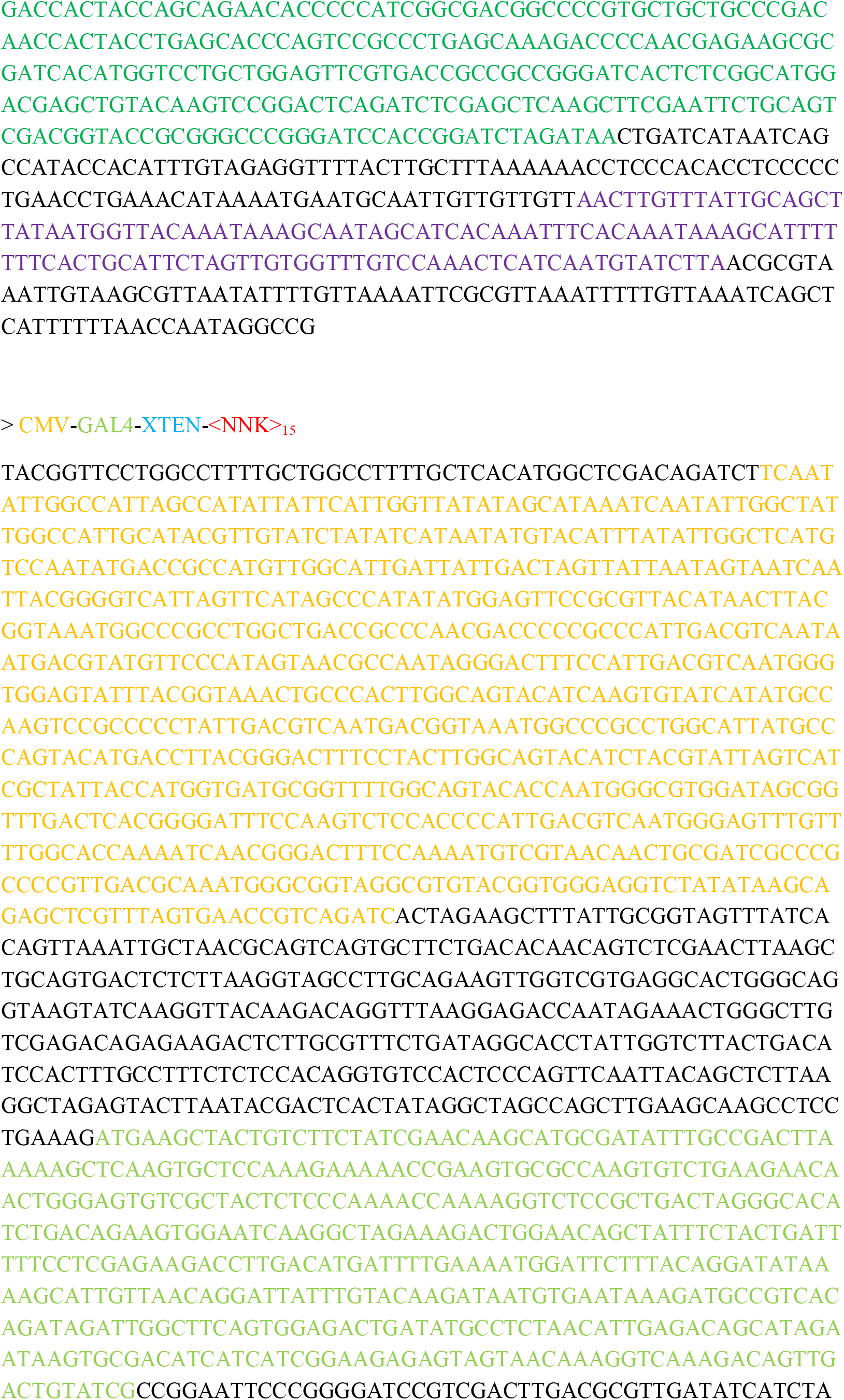

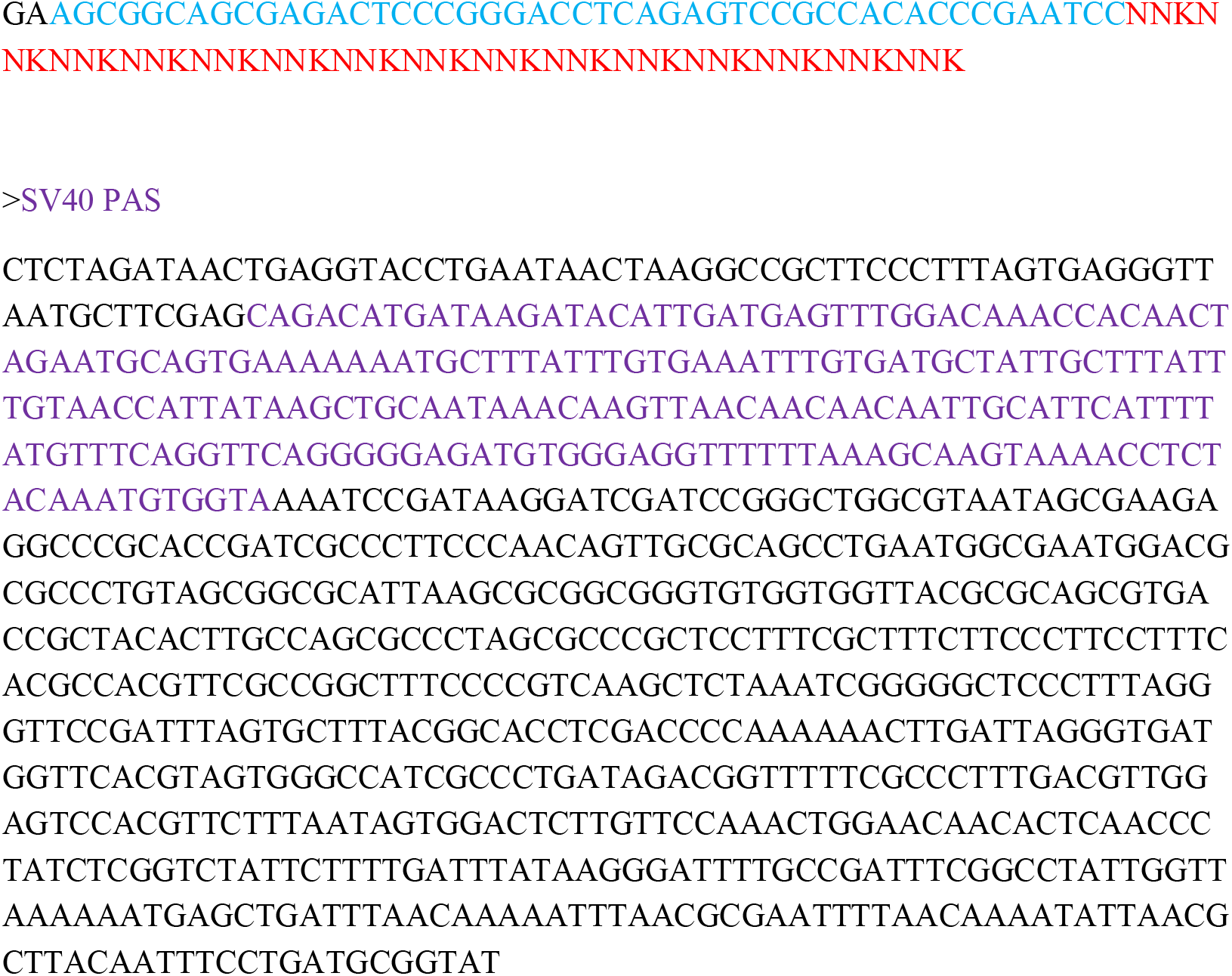

